# RecAlign: A* recombination-aware sequence to graph mapping

**DOI:** 10.1101/2025.01.18.633308

**Authors:** Paola Bonizzoni, Davide Cesare Monti, Gianluca Della Vedova, Brian Riccardi, Raffaella Rizzi, Jouni Siren

## Abstract

Pangenomics and long reads bring the promise of integrating read mapping with variant calling, since a pangenome encodes a reference genome that incorporates evolutionary or population aspects, while even a single long read can provide a good evidence of different kinds of variants (not only the single nucleotide variants that can be easily observed by short reads). This promise needs to be fulfilled by the development of new read mapping approaches that are tailored for that purpose. This paper focuses on integrating recombination events, that are key in bacteria, into read mapping. A first approach in that direction [ACBC^+^24] provides an exact dynamic programming algorithm that is too slow to manage multiple recombinations or long reads. We present a novel A* algorithm for recombination-aware sequence-to-graph mapping that significantly reduces running time by incorporating haplotype information and an efficient heuristic function. Our tool, RecAlign, demonstrates up to a two-order magnitude improvement in time and space complexity over [ACBC^+^24] and efficiently handles multiple recombinations.

## 1 Introduction

Sequence-to-graph alignment is a fundamental task in bioinformatics for sequence comparison against a pangenome graph [B^+^22]. A pangenome graph is a recent paradigm adopted to represent a reference genome [LAE^+^23] and it will become in the near feature the standard for representing collections of genomes in a population. Together with the introduction of pangenome graphs, it has been observed that distinguishing haplotype information is crucial [SGN^+^19] in dealing with such reference graphs. Indeed, given a collection of genomes, haplotypes express the evolution of the species and, among the guiding events, recombinations that occur at a different frequency rates in species, play a crucial role in relating individual haplotypes. For example, in bacterial pangenomes [DM10] recombinations are a relevant phenomenon and tools for detecting them are extremely useful in reconstructing the evolutionary history. In [ACBC^+^24] a recombination-aware sequence-to graph alignment approach has been proposed to directly handle recombination events in a pangenome graph. Indeed, a pangenome graph is meant to be at least as informative as a multiple sequence alignment, and recombinations have traditionally been detected through multiple sequence comparison [SRS02]. While [ACBC^+^24] is the first algorithmic approach that incorporates the notion of recombination into a sequence-to-graph alignment, its implementation manages effectively at most one recombination, since allowing multiple recombinations increases the computational complexity, which is already linear in the *product* between the size of the query and the size of the graph — this makes the approach too expensive when dealing with entire genomes and large pangenome graphs. Indeed, unless the Strong Exponential Time Hypothesis (SETH) is false, as proved in [EMTG23], a subquadratic algorithm cannot be achieved even in the case of restricted classes of graphs, thus encouraging the investigation of algorithmic approaches that are fast enough in practice and on average. Notice that adding recombinations to the alignment makes sense only if we distinguish paths of the graph representing distinct haplotypes. This information is not considered by some fast sequence-to-graph aligners, such as [RM20], which are able to align even long reads but disregards haplotype information and uses heuristics to speed up the average case.

In this paper, we greatly reduce the running time of RecGraph[ACBC^+^24], allowing it to compute a read-to-graph mapping that can have multiple recombinations. While RecGraph‘s dynamic programming approach allows *k* recombinations, its time complexity is exponential in *k*, therefore it is impractical on viral genomes or long reads even when *k* = 2. We design an A* approach that successfully manages haplotype information and represents recombinations as switches between distinct haplotype paths of the graph, greatly reducing the running time of recombination-aware sequence-to-graph mapping. While the A* approach has been successfully applied to speed-up pairwise sequence alignment [GKI24] and sequence-to-graph alignment [IBV22], those approaches do not consider recombinations. Our main technical contribution is a new, recombination-aware, heuristic function *h* (1) fits into the RecGraph approach and (2) is a good lower bound on the value of a recombination-aware sequence-to-graph alignment between a suffix of the query string and a portion of the pangenome graph. Since our function *h* provides a lower bound of the optimum, the A* properties guarantee that we obtain an optimal alignment, while exploring only a small portion of the alignment graph — notice that RecGraph explores the entire alignment graph, which is the main reason of its slower running times. In addition, we change the representation of a recombination event from *recombination edge*, as described in [ACBC^+^24], to *switch node*, thus reducing the size of feasible solution spaces, further increasing the performance. The algorithmic approach has been implemented and tested in the tool named RecAlign to test the performance of the new approach, showing that we can achieve a decrease in time and space by up to two orders of magnitude compared to RecGraph. Most importantly, RecAlign can efficiently handle multiple recombinations and is as accurate as RecGraph in detecting the paths involved in the recombinations.

## 2 Preliminaries

A string is a sequence *s* = *s*[1] … *s*[*n*] of symbols drawn from an alphabet Σ. We denote by |*s*| the length of *s*. We will use the bracket notation to denote substrings of *s*, that is, given *i, j* with 1 ≤ *I* ≤ *j* ≤ |*s*|, we denote by *s*[*i* : *j*] the substring *s*[*i*] … *s*[*j*]. A prefix (resp. suffix) of *s* ending (resp. starting) at position *i* will be denoted by *s*[: *i*] (resp. *s*[*i* :]).

In this paper we consider *canonical Variation Graphs* [B^+^22], which are directed graphs representing a set of haplotypes, where each vertex is labeled by a character. The algorithm we propose will explore an alignment graph built from the paths of the input variation graph; this means that pangenome graphs are not required to be acyclic, unlike [ACBC^+^24].

### Definition 2.1.

(Canonical Variation Graph) *A* canonical variation graph *is a tuple G* = ⟨*V, A, P, λ*⟩ *where V is a set of* vertices, *A* ⊆ *V* × *V is a set of* directed arcs, *P is a set of distinguished paths called* haplotypes, *and λ* : *V* → Σ *is the* labelling function.

We assume, without loss of generality, that *V* contains a unique source *s* and sink *t*, both labeled by the empty string, such that each haplotype *p* ∈ *P* starts in *s* and ends in *t*. Furthermore, we assume that each arc (*u, v*) ∈ *A* is traversed by some *p* ∈ *P*.

We extend the labelling function from vertices to paths, by letting *λ*(*π*) = *λ*(*v*_1_)*λ*(*v*_2_) … *λ*(*v*_*k*_), for any path *π* = ⟨*v*_1_, *v*_2_, …, *v*_*k*_⟩ in *G*.

An *alignment* of a string *s* of length *n* against a path *π* = ⟨*v*_1_, *v*_2_, …, *v*_*k*_⟩ of *G* is a sequence ⟨(*x*_*i*_, *y*_*i*_)⟩ of *q* ordered pairs such that *x*_*i*_ ∈ {1, …, *k*, −}, *y*_*i*_ ∈ {1, …, *n*, −} satisfying the following: (i) for any 1 ≤ *i < j* ≤ *q*, if − ∈*/* {*x*_*i*_, *x*_*j*_} then *x*_*i*_ *< x*_*j*_, (ii) for any 1 ≤ *i < j* ≤ *q*, if − ∈*/* {*y*_*i*_, *y*_*j*_} then *y*_*i*_ *< y*_*j*_, (iii) for any 1 ≤ *i* ≤ *q*, at least one of *x*_*i*_, *y*_*i*_ is not −, (iv) {1, …, *k*} = {*x*_*i*_ : *x*_*i*_ *≠*−}, and (v) {1, …, *n*} = {*y*_*i*_ : *y*_*i*_ *≠*−}.

If the indel − ∈*/* {*x*_*i*_, *y*_*i*_}, then the pair (*x*_*i*_, *y*_*i*_) represents a *match* if *λ*(*v*_*x*_*i*) = *s*[*y*_*i*_], otherwise it represents a *mismatch* (or *substitution*). If the indel − ∈ {*x*_*i*_, *y*_*i*_}, then the pair (*x*_*i*_, *y*_*i*_) represents an *indel* (*i*.*e*., an *insertion* either in *π* or in *s*). Notice that this definition essentially models the standard sequence alignment between *s* and the label of the path *π*.

The value of an alignment is defined as the sum of the values *δ*(*x*_*i*_, *y*_*i*_) for each 1≤ *i* ≤ *q*, where *δ* is a *scoring scheme* function, assigning a value to each pair of the alignment. Please note that *δ* assigns positive values if the goal is to compute an alignment of minimum cost.

### 2.1 Computing recombination-aware alignments

A node *v*, different from the source *s* and the sink *t*, that belongs to two distinct haplotypes in *P* will be called a *switch node*. Moreover, we call *π* = ⟨*v*_1_, *v*_2_, …, *v*_*k*_⟩ an *r-switch path* a if there exist *r* distinguished switch nodes 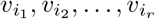 (with 1 ≤ *i*_1_ *< i*_2_ *<* … *< i*_*r*_ ≤ *k*) such that: (i) if 1 *< p < r*, then 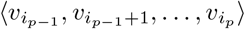 and 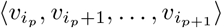 are subpaths of the two different haplotypes, (ii) if *i*_1_ *>* 1, then 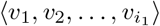 and 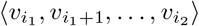 are subpaths of the two different haplotypes (iii) if *i*_*r*_ *< k*, then 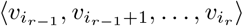 and 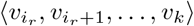 are subpaths of the two different haplotypes. Observe that an *r*-switch path represents a path of *G* = ⟨*V, A, P, λ* ⟩ (but not necessarily a path in *P*) with *r* recombinations; in particular, a haplotype in *P* is a source-sink *0-switch* path.

Finally, we introduce a constant *ρ* which is the recombination cost. We are now ready to define the problem we want to solve.

#### Definition 2.2.

(Recombination-aware Alignment Problem (RAAP)) *Let G* = ⟨*V, A, P, λ*⟩ *be a canonical variation graph, let Q be a* query sequence, *δ be a scoring scheme, ρ be a recombination cost, and let R* ≥ 0 *be an integer. The* Recombination-aware Alignment Problem *(RAAP) asks for a minimum cost alignment (w*.*r*.*t. δ and ρ) of Q against all r-switch paths of G, with r* ≤ *R*.

A dynamic programming solution for RAAP has been described in [B^+^22] and is reported here. We define *M* [*v, i, p, r*] as the minimum cost of an alignment of *Q*[: *i*] against any *r*_1_-switch path *z* of *G* = ⟨*V, A, P, λ*⟩, with *r*_1_ ≤ *r*, where *z* starts with the vertex *s* and ends with the vertex *v* of the haplotype *p* ∈ *P*. Thus, the optimal solution of an instance of RAAP is given by min_*p∈P*_ {*M* [*t*, |*Q* |, *p, R*]}, where *t* is the sink node of the canonical variation graph *G*.

A recursive definition of *M*, where *v* ≠ *s*, (*u, v*) is the arc of *p* incoming into *v*, and *i >* 0, is the following:

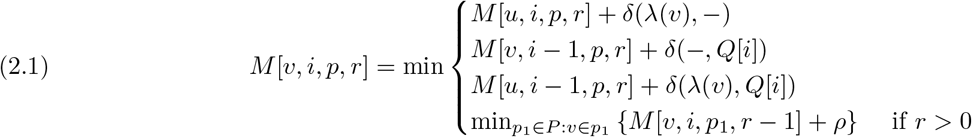

The first three cases of Equation (2.1) are the *fixing* moves, *i*.*e*. those for which the alignment continues along the haplotype *p*, while the last case represents the *recombination* moves, *i*.*e*. those for which the alignment switches from *p*_1_ to *p*.

## 3 Solving RAAP via the A* algorithm

In this section we describe how we can use the A* approach [HNR68], which is a technique that speeds up Dijkstra’s algorithm [Dij59] to compute a source-to-target shortest path in a weighted directed graph, to reduce the time to compute an alignment. Both the original Dijkstra’s algorithm and A* maintain two disjoint sets of vertices of the graph; the close set *C*, containing the vertices *v* ∈ *V* for which the source-to-*v* distance *d*(*v*) has been evaluated, and the frontier *F*, that is the vertices that have not been evaluated but are reachable from *C* by traversing a single arc. Initially, the close set contains only the source, the frontiers all vertices that are adjacent to the source, and at each iteration of the Dijkstra’s algorithm the vertex of the frontier that is closer to the source is added to the close set, until the sink is reached.

The A* instead selects the vertex in the frontier based on a heuristic function *h* : *V* → ℚ which provides an estimate of the minimum distance from *v* to the target: the better the estimate, the smaller the close set and the frontier. More precisely, we select (and move to the close set *C*) the vertex *v* of the frontier minimizing *f* (*v*) = *g*(*v*) + *h*(*v*), where *g*(*v*) is an upper bound on the distance *d*(*v*) from the source to *v*. Moreover, we update the upper bound *g*(*u*) of each neighbor *u* of *v* that does not belong to the close set.

The choice of the heuristic function *h* is crucial. In fact, if *h* is *admissible*, that is *h*(*v*) is never larger than the cost of the shortest path from *v* to the target node, then the optimal solution is always found. On the other hand, if *h* is admissible but too conservative, then a large portion of the graph will be explored, rendering A* ineffective. Notice that when *h* = 0, then A* is precisely Dijkstra’s algorithm.

Our goal is to use A* to compute *M* [*t*, |*Q* |, *P, R*] and find the optimal alignment of the query sequence *Q* against *G* with at most *R* recombinations. Our approach is inspired from [Gus97, GKI24] for the encoding of the recurrence through a directed graph, referred to as *alignment graph*, and exploits the A* algorithm in order to efficiently compute the shortest paths leading to the solution. The following sections describe the alignment graph encoding our recurrence, the estimate function *h* we use for A*, and how we explore the *alignment graph* in order to find an optimal alignment.

In what follows, the scoring scheme *δ* will be the classic edit distance score (*i*.*e*. mismatches, insertions and deletions at value 1, matches at value 0).

### 3.1 Alignment Graph

Considering the recurrence Equation (2.1), we say that entry *û* = (*u, j, p*^*′*^, *r*^*′*^) *yields* entry 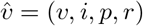 whenever *M* [*û*] appears in the recursive definition of 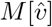. Furthermore, the *cost of yielding* 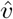 *from û* is *δ*(·) (if *p* = *p*^*′*^) or *ρ* (otherwise). The *alignment graph* encoding the recurrence (Equation (2.1)) is a weighted graph *G*_*A*_ = ⟨*V*_*A*_, *E*_*A*_, *W*_*A*_⟩ such that:

1. *V*_*A*_ is the set of entries of *M*, plus two extra vertices *ŝ* and 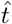 (respectively, the source and the destination of *G*_*A*_);
2. for every pair of entries *û*, 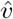 of *M*, there is an edge (*û*, 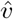) ∈ *E*_*A*_ whenever *û* yields 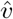. Furthermore, there are edges (*ŝ*, (*s*, 0, *p, r*)) and (*f*_*p*_, |*Q*|, *p, R*), 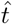) for every *p* ∈ *P* (recall that *f*_*p*_ is the last vertex of haplotype *p* before reaching *t* in *G*);
3. for every arc (*û*,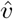) ∈ *E*_*A*_, the cost *W*_*A*_(*û*, 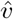) is the cost of yielding 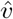 from *û* if both are entries of *M*, otherwise it is 0 if either *û* = *ŝ* or 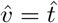.

As already stated, the optimal (minimum) value of the alignment of *Q* against *G* with at most *R* recombinations is given by *M* [*t, Q, P, R*] = min_*p∈P*_ {*M* [*f*_*p*_, |*Q* |, *p, R*]}. It is immediate to see that such value is the shortest distance from *ŝ* to 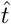 in *G*_*A*_. Thus, we have reframed the *RAAP* problem to the shortest-path problem.

### 3.2 Estimate functions

In this section, we are going to present two different estimate functions used by our approach, respectively *h*_*s*_ and *h*_*c*_. The first uses all matches between the query *Q* and the graph *G*, while the second uses only matches that can be chained together.

#### 3.2.1 Seed Estimate Function *h*_*s*_

The estimate function *h*_*s*_ : *V*_*A*_ → ℝ we propose is inspired by [GKI24] and is specialized for sequence-to-graph alignment in the presence of recombinations. Recall that, given a vertex *v*_*A*_ = (*v, i, p, r*) of the alignment graph, the value *h*(*v*_*A*_) must be a lower bound of the shortest distance (in *G*_*A*_) from *v*_*A*_ to the target 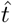. Notice that each path in the alignment graph from *v*_*A*_ to 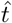 corresponds to an alignment of *Q*[*i* + 1 :] and a portion of the graph *G* where the source is *v* instead of *s*.

Let us consider a path in the alignment graph from *v*_*A*_ to 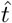, we can divide such a path into portions; some of them correspond to errorless matches between the query string and the haplotype, and some contain at least a mismatch or a recombination switch. The alignment cost is at least as large as the number of portions containing mismatches or switches.

In what follows, we will give precise definitions and constructions for both the zero-recombination case and the general case with multiple recombinations allowed. The careful reader should have already understood that the estimate for vertex (*v, i, p, r*) will depend only on *p, i* and thus, in the subsequent sections, we will refer to the estimate as *h*(*p, i*).

##### Seeding and matching

For simplicity of exposition, we assume that *Q*[*i* : *j*] = *Q*[*i* :] when *j >* |*Q*|, that is when the end position of a substring is beyond the boundary of the string. We will split the query string *Q* into non-overlapping substrings of constant length, the *seeds*, and we consider such seeds as “macro-symbols” to match against the variation graph. More formally a seed is defined as follows:

###### Definition 3.1.

*Let Q* ∈ Σ^*n*^ *be a* query string, *and k (*1 ≤ *k* ≤ *n) be the* seed length. *Then, a k*-seed of *Q is a pair* ⟨*i, Q*[(*i* − 1)*k* + 1 : *ik*]⟩ *for* 1 ≤ *i* ≤ ⌊*n/k*⌋.

Notice that a *k*-seed is uniquely identified by its index *i*, therefore we can use *QS*_*k*_(*i*) to denote the string *Q*[(*i* − 1)*k* + 1 : *ik*]. Now, we introduce the notion of matching of a *k*-seed.

###### Definition 3.2.

*Let G* = ⟨*V, A, P, λ*⟩ *be a variation graph, p* ∈ *P a distinguished path of G and Q a query string. An* exact (*k, p, Q*)-match *(or simply a match, when there is no ambiguity) is a pair* ⟨*u, i*⟩ *s*.*t*.:

1. 1 ≤ *u* ≤ |*λ*(*p*)|,
2. 1 ≤ *i* ≤ ⌊*n/k*⌋,
3. *λ*(*p*)[*u* : *u* + *k* − 1] = *QS*(*i*).

A (*k, p, Q*)-match ⟨*u, i*⟩ will be also indicated with ⟨*i, j*⟩_*p*_ to remark the fact that it refers to haplotype *p*. Moreover, we denote by *M*_*k*_(*p, Q*) the set of all exact (*k, p, Q*)-matches of the *k*-seeds of *Q* with respect to haplotype *p*. In our implementation, to compute the various sets *M*_*k*_(*p, Q*) for each *p* ∈ *P* we build an *Lt-Fm-Index* [AW21] for each path^1^.

##### Definition of the estimate function *h*_*s*_

As already anticipated in Section 3.2, the estimate function *h*_*s*_ used by our approach is based on the matches between the seeds of *Q* and each haplotype.

The main idea is that, given an index *i*, every seed with index *j > i* that does not have a corresponding match ⟨·, *j*⟩ ∈ *M*_*k*_(*p, Q*) will require at least an edit operation in the alignment between *p* and *Q*[*ik* :]. For this reason, if we define *ed* = min{*δ*(*σ*_1_, −), *δ*(−, *σ*_1_), *δ*(*σ*_1_, *σ*_2_) | *σ*_1_, *σ*_2_ ∈ Σ, *σ*_1_ ≠ *σ*_2_}, then the cost of the alignment between *p* and *Q* will increase, at least, by *ed* for each seed without a match in *C*_*k*_(*p, Q*).

We can now give a formal definition of the estimate function *h*_*s*_.

###### Definition 3.3.

(*h*_*s*_) *Let M*_*k*_(*p, Q*) *be the set of matches between Q and a path p* ∈ *P*. *The estimate function h*_*s*_ : (*P* × *S*_*k*_(*Q*)) → ℕ *as:*

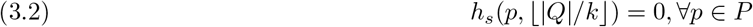

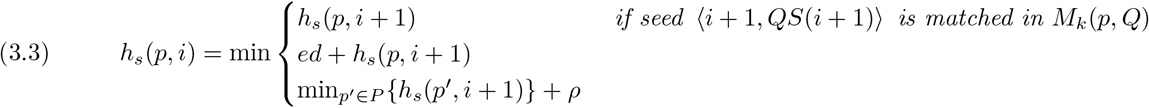

The first two cases of Equation (3.3) correspond to alignments that do not have a recombination, while the last case corresponds to alignment with a recombination. Note that this estimate allows an unbounded number of recombination events.

#### 3.2.2 Chain Estimate Function

*h*_*c*_ We are also providing a second estimate function used by RecAlign, based on the seed estimate function but with the addition of a chaining step. The idea is that, once all *M*_*k*_(*p, Q*) have been calculated, we want to extract chains of matches for each haplotype that can form the backbone of an alignment. First we give a partial order on matches, then we will provide a definition of chain respecting such partial order.

##### Definition 3.4.

*Let* ⟨*u*_1_, *i*_1_⟩_*p*_ *and* ⟨*u*_2_, *i*_2_⟩_*p*_ *be two (k, p, Q)-matches with respect to the same haplotype p. Then*, ⟨*u*_1_, *i*_1_⟩_*p*_ ≺ ⟨*u*_2_, *i*_2_⟩_*p*_ *if and only if u*_1_ + *k < u*_2_ *and i*_1_ *< i*_2_.

A *chain* of *M*_*k*_(*p, Q*) is a (≺)-increasing sequence ⟨⟨*u*_1_, *i*_1_⟩, …, ⟨*u*_*z*_, *i*_*z*_⟩⟩ of matches in *M*_*k*_(*p, Q*), that is ⟨*u*_*h*_, *i*_*h*_⟩ ≺ ⟨*u*_*h*+1_, *i*_*h*+1_⟩ for each *h*.

Moreover, we are interested in finding the *seed longest chain*, denoted by *C*_*k*_(*p, Q*[*ik* :]) for every path *p* ∈ *P* and seed in *Q*, defined as follows:

##### Definition 3.5.

(Seed Longest Chain) *Let M*_*k*_(*p, Q*) *be a set of* (*k, p, Q*)*-matches, then a chain* ⟨⟨*u*_1_, *i*_1_⟩, …, ⟨*u*_*z*_, *i*_*z*_⟩⟩ *in M*_*k*_(*p, Q*) *is a* seed longest chain *C*_*k*_(*p, Q*[*ik* :]) *if and only if:*

1. *(seed) there is no match* (*u, j*) *in C*_*k*_(*p, Q*[*ik* :]) *such that j < i, and*
2. *(longest) no other chain of M*_*k*_(*p, Q*) *contains more matches*.

We compute a seed longest chain by a simple variant of the well-known algorithm for the construction of a *Longest Increasing Subsequence* [Fre75], where the order used prioritize the position of the match on the query string over the position on the haplotype *p*.

##### Definition of the estimate function *h*_*c*_

Every chain *C*_*k*_(*p, Q*[*ik* :]) can be seen as the largest subset of matches from *M*_*k*_(*p, Q*), with seeds greater than *i*, that can be chained together. Each seed that is not in *C*_*k*_(*p, Q*[*ik* :]) incurs a cost, but we have to determine if such a cost is due to a mismatch or to a recombination. To that purpose, we compute the *chain costs* for every seed *i*, containing the paths with minimum cost for every path *p*, using only the matches present in the chain *C*_*k*_(*p*, [*ik* :]). Notice that the *h*_*s*_ heuristic does not incur in a cost whenever a seed matches anywhere in the haplotype, that is even if two matches cannot be chained.

###### Definition 3.6.

(Chain Costs) *Let Chains*_*i*_ = {*C*_*k*_(*p, Q*[*ik* :]) | *p* ∈ *P*} *be the set of seed longest chain for every path p* ∈ *P, we define the chain costs* 𝒞_*i*_ *of size* |*P* | × ⌊|*Q*|*/k*⌋ *as:*

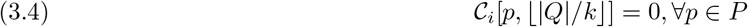

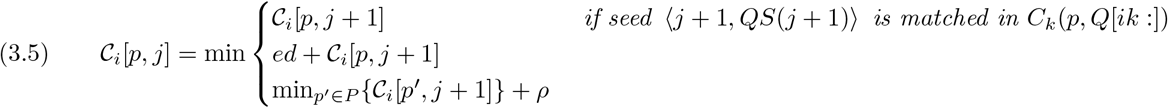

After having computed the chains cost matrices for every seed, the column corresponding to the seed from which the chain was built will contain the minimum cost of the alignment from that position to the target. Or, more formally, we can define the estimate function used by our approach as:

###### Definition 3.7.

(Estimate Function) *Let* C_*i*_ *be the chains cost matrix for the i-th seed of Q, then we define the estimate function h* : (*P* × *S*_*k*_(*Q*)) → ℕ *as:*

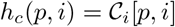

#### 3.2.3 Proof of optimality

In order to prove that RecAlign is able to compute the optimal solution for RAAP, we have to show that our estimate function *h*_*s*_ is actually a lower bound on the cost of the path from a generic vertex (*v, j, p, r*) to 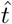 in *G*_*A*_.

##### Theorem 3.1.

*Let* (*v, j, p, r*) *be a vertex of the alignment graph G*_*A*_, *then h*_*s*_(*p*, ⌊*j/k* ⌋) *is less than or equal to the distance from* (*v, j, p, r*) *to* 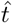 *in G*_*A*_.

*Proof*. Let *v*_*a*_ = (*v, j, p, r*), let *path*(*v*_*a*_) be a shortest path from *v*_*a*_ to the target node 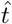 in *G*_*A*_, and let 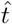 be the length of such path. Moreover, let *i* = ⌊*j/k*⌋. Notice that each seed ⟨*i, QS*(*i*) ⟩ identifies the subpath *path*(*v*_*a*_, *i*) of 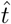 starting with the first vertex of the form (·, (*i* − 1)*k* + 1, ·, ·) and ending with the last vertex of the form (·, *ik*, ·, ·). All such subpaths are vertex-disjoint and correspond to the portion of the optimal alignment involving the substring *QS*(*i*). W.l.o.g. we can assume that *ρ > ed*. If *j >* ⌊*n/k*⌋ *k*, then *h*(*p, i*) = 0 by definition, therefore 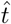 and we need to consider only the case *j* ≤ ⌊*n/k*⌋ *k*, that is the position *j* belongs to the seed with index *i*.

We only need to consider cases of the form *v*_*a*_ = (*v*, (*i* + 1)*k*−1, *p, r*), that is involving the last character of a seed. In fact, for any other *j*, the optimal path from (*w, j, p, r*) to the target vertex includes the optimal path from (*v*, (*i* + 1)*k* − 1, *p*_1_, *r*_1_), for some *v, p*_*q*_, *r*_1_, to the target vertex, hence 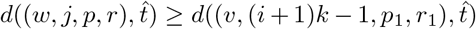, and 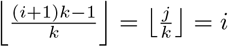. Moreover, we only need to consider cases where |*Q*| mod *k* = 0, as the heuristic will not change for the trailing symbols of *Q* that can not be included in a seed.

Now, suppose that 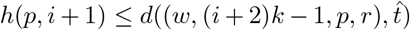, ∀*p*∈ *P*. Because of Definition 3.7, *h*(*p, i*) can take the minimum between three values:

1. *h*(*p, i* + 1)
2. *h*(*p, i* + 1) + *ed*
3. *h*(*p*^*′*^, *i* + 1) + *ρ*

If we are in case 1, it is immediate to see that, since 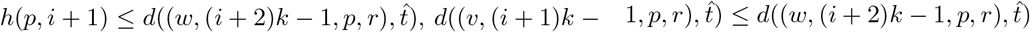 and *h*(*p, i*) = *h*(*p, i* + 1), then 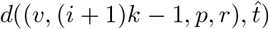.

If we are in case 2, instead, we can see the distance 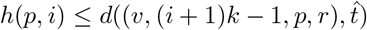 as composed of two parts: the distance *d*\* = *d*((*v*, (*i* + 1)*k* − 1, *p, r*), (*w*, (*i* + 2)*k* − 1, *p, r*)) plus 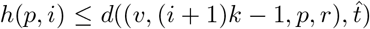, where *d*\* represents the cost of aligning *Q*[(*i* + 1)*k* : (*i* + 2)*k* − 1] against *p*[*v* : *w*]. In particular, if at least one edit operation is required in this alignment, then, since it will cost at least *ed*, we can see that 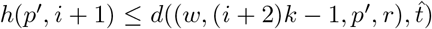. For this reason, we have to consider only the case where *d*\* = 0. However, this implies that *Q*[(*i* + 1)*k* : (*i* + 2)*k* − 1] and *p*[*v* : *w*] are perfectly matched. Please note that *Q*[(*i* + 1)*k* : (*i* + 2)*k* − 1] is actually the (*i* + 1)-th seed and, since we are in case 2, it is unmatched in *M*_*k*_(*p, Q*) even if we should have a match *m* = ⟨*v, i* + 1⟩ because it can be aligned with 0 cost. This means that is impossible to have *d*\* = 0 and *m ∉ M*_*k*_(*p, Q*).

Lastly, we have to consider case 3. In particular, we have two possible outcomes: we have a recombination in the shortest path covering the positions *Q*[(*i* + 1)*k* : (*i* + 2)*k* − 1] or there is no recombination over those positions. If there is no recombination, since *h*(*p*^*′*^, *i* + 1) + *ρ* ≤ *h*(*p, i* + 1) + *ed*, we can use what we prove for case 2, and this implies that 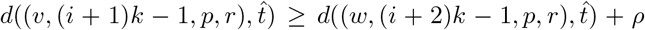. If there is at least a recombination, instead, by hypothesis we already know that 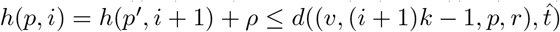, and, since at least one recombination occurred, 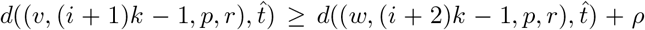. For this reason, we can also affirm that 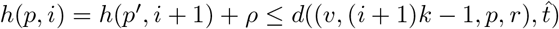.

### 3.3 A* exploration

In this section, we describe the actual A* exploration on *G*_*A*_ and how to obtain the shortest path from source to target node, after determining chains *C*_*k*_(*p, Q*) and the estimate function *h*(*p, i*). We are able to do that by keeping updated two sets: a *min-priority queue F* (the frontier), which represents the nodes to be expanded with A*, and a *hash-set E*, the subgraph of *G*_*A*_ containing the explored positions. Please note that *G*_*A*_ can be inferred starting from *G* and *Q*, for this reason, there is no need to initialize it *a priori*, but it is built during this step, with only the nodes actually explored, using *E*.

**Initialization** Firstly, the source node *ŝ* is added both to *F* and *E*. Then, ∀*p* ∈ *P*, we initialize the starting vertex of the path *v*_*s,p*_ = (0, 0, *p*, 0), with priority equal to 0 + *h*(*p*, 0), parent equal to *ŝ* and *g* = 0. We then start a classic A* exploration of the alignment graph.

**Skip ahead step** When the node selected by the A* procedure *n*_*min*_ = (*v, i, p, r*) coincides with the starting position of a match included in the chain *C*_*k*_(*p, Q*[*ik* :]), it is possible to skip all nodes until the end of the match is reached. Formally:

#### Definition 3.8.

*Let C*_*k*_(*p, Q*[*ik* :]) *be a length-maximum chain in M*_*k*_(*p, Q*), *and n*_*min*_ = (*v, i, p, r*) *be the current vertex extracted from F*. *If* ⟨*v, ik*⟩_*p*_ ∈ *C*_*k*_(*p, Q*[*ik* :]), *then the next position to be explored is* (*v* + *k, i* + *k, p, r*).

Since Theorem 3.1 shows that *h* is admissible, once we reach the vertex *v* ∈ *G*_*A*_, we also have the shortest path from the source *v*_*a*_ to *v* — all other paths from *v*_*a*_ to *v* are not relevant and are not explored. In order to penalize the exploration of the same portions of the alignment graph, without improving the alignment, this match is removed from the chain and the score of *h*(*p, i*) is increased by one for each position *i* preceding this match.

**Termination** The exploration of *G*_*A*_ stops as soon as a *target* node is extracted from *F*, this means that the procedure stops the first time a node (*t*, |*Q*|, *p, r*) is expanded.

#### 3.3.1 Differences in semiglobal alignment

The procedure described in Section 3.3 describes a global alignment, where we are interested in aligning the whole query *Q* against the whole paths in *G* = (*V, A, P, l*). If we want to obtain a semiglobal alignment, where we are interested in aligning the whole *Q* against a subgraph of *G*, we have to change the initialization phase to include all arcs (*v*_*a*_, (*v*, 0, *p*, 0)) and all arcs ((*v*, |*Q*|, *p, R*), *v*_*ω*_). All such additional arcs have weight zero. Notice also that the implementation does not necessarily store explicitly those arcs.

## 4 Results

We implemented the method described in Section 3 in Rustand the tool, named RecAlign is available at https://github.com/AlgoLab/RecGraph on the branch a starunder the MIT license. RecAlign takes as input a graph in GFA format and a set of sequences in FASTA format and produces as output the optimal alignment of each sequence against the graph in the GAF format.

We designed an experimental evaluation in order to test our current implementation against the old RecGraph and other sequence to graph aligners, namely GraphAligner [RM20] and Minichain [CGJ24]. Observe that a comparison of RecGraph with other non haplotype-aware tools have also been presented in [ACBC^+^24].

### 4.1 Experiment 1

In this experiment, we replicated Experiment 2 of [ACBC^+^24] to evaluate the quality of RecAlign in predicting recombinations. We built a pangenome graph using 11 variants of the *slpA* gene of *C. Difficile* [DDA^+^13], and then we simulated 994 new genes, each of which might be a recombinant of two random variants, with a randomly generated breakpoint. Please note that in order to build the graph we used pggb[GG23] for RecAlign, since RecAlign is able to handle graph with cycles and inversions, while the graph used by RecGraph was acyclic. Lastly, we aligned each simulated sequence against the graph using both the new *A\** version and the old one. The goal was to determine if RecAlign was actually able to match RecGraph’s precision in detecting recombinations.

Considering the results obtained, in each simulated sequence without a recombination (*n*= 102) RecAlign correctly computed an alignment that does not include a recombination, and, for each sequence with a recombination (*n* = 892), RecAlign produced an alignment with the correct recombination and paths interested in the alignment. However, considering the location of the predicted breakpoint, RecAlign obtained an inferior result compared to RecGraph, with a mean distance of 17 bp, compared to the 1.64 bp distance of the old version.Please note that these results are expected, since RecAlign considers recombination as *switches* while RecGraph was able to add a *recombination edge*, thus improving the precision of the position of the recombination event. However, since the simulated sequences were approximately 2200 bp long, this lower precision is not really remarkable, and it is justified by the improvements in performances and the possibility of identifying multiple recombinations that we are going to show in the next experiment.

### 4.2 Experiment 2

In this experiment, the aim was to show the improvements in performances of our approach. For this reason, we use the data available at https://github.com/ekg/HLA-zoo which are usually considered among the most difficult regions for an aligner. The repository contains 28 genes of the HLA complex, with multiple variants, extracted from the human genome reference GRCh38. In humans, the HLA (Human Leukocyte Antigen) genes encode proteins that play a crucial role in the immune system by presenting peptide antigens to T cells and these genes are known for their extreme diversity, with numerous distinct alleles. We chose this particular dataset because we wanted to test RecAlign capabilities on longer reads and on a series of genes with high differences.

We downloaded the variation graphs from https://github.com/ekg/HLA-zoo and used them to generate sequences representing whole genomes, introducing errors with different rates (0,3,5 and 10%) by randomly changing, deleting or duplicating bases of the original haplotype. We have also introduced some recombination events (at most 2 for each sequence) by mixing different haplotypes. Then we aligned each simulated sequence against its respective graph. We chose to use pangenome graphs produced by spoa(https://github.com/rvaser/spoa), a tool that is based on the POA algorithm [LGS02] with the addition of SIMD instructions. We chosethem in order to have graph without cycles, that are not correctly handled by RecGraph and Minichain.

Since the simulated sequences have lengths that span from around 3kbp up to around 18kpb, we were not able to successfully produce any alignments for the longer ones with RecGraph in a reasonable time. We then decide to select 3 of the shortest gene, in order to compare RecAlign against RecGraph, and we used only RecAlign for the complete set of all genes available in https://github.com/ekg/HLA-zoo.

#### 4.2.1 Comparison with RecGraph

In this part of the experiment, we selected 3 of the shortest genes (K-3138, H-3136 and V-352962) and we simulated read with increasing error rates but without recombination events, and reads without errors but presenting 2 recombination events. The goals of this experiment were to compare the differences in RecAlign and RecGraph, and determine if the performances of RecAlign were affected more by the error rates or by the presence of recombination events.

Firstly, we measured the edit distances between each sequence and the portion of the graph identified by the alignment, the results are in figure 1. RecAlign was able to match the performances of RecGraph in every genes, showing slight improvements when considering reads presenting 2 recombination events, since RecGraph is able to identify at most one recombination event. We then measured the performances of the two tools, considering time and memory required during the alignment procedure, the results are reported in figures 2 and 3. These results highlight an improvement in terms of both speed and occupied memory by a factor of at least 10^2^. In particular, we can note that while RecGraph is not affected by the error rate of the sequences, RecAlign is largely affected. Both memory and time required by the alignments increase with the error rate, but even with an error rate of 10% (that is even higher than the error rate of current sequencing technologies) we were largely better than RecGraph. On the contrary, the experiment suggests that the number of recombination present in the sequences is not affecting the performances of RecAlign, in fact we can see that both time and memory are quite similar to the case without errors and without recombinations.

**Figure 1:**
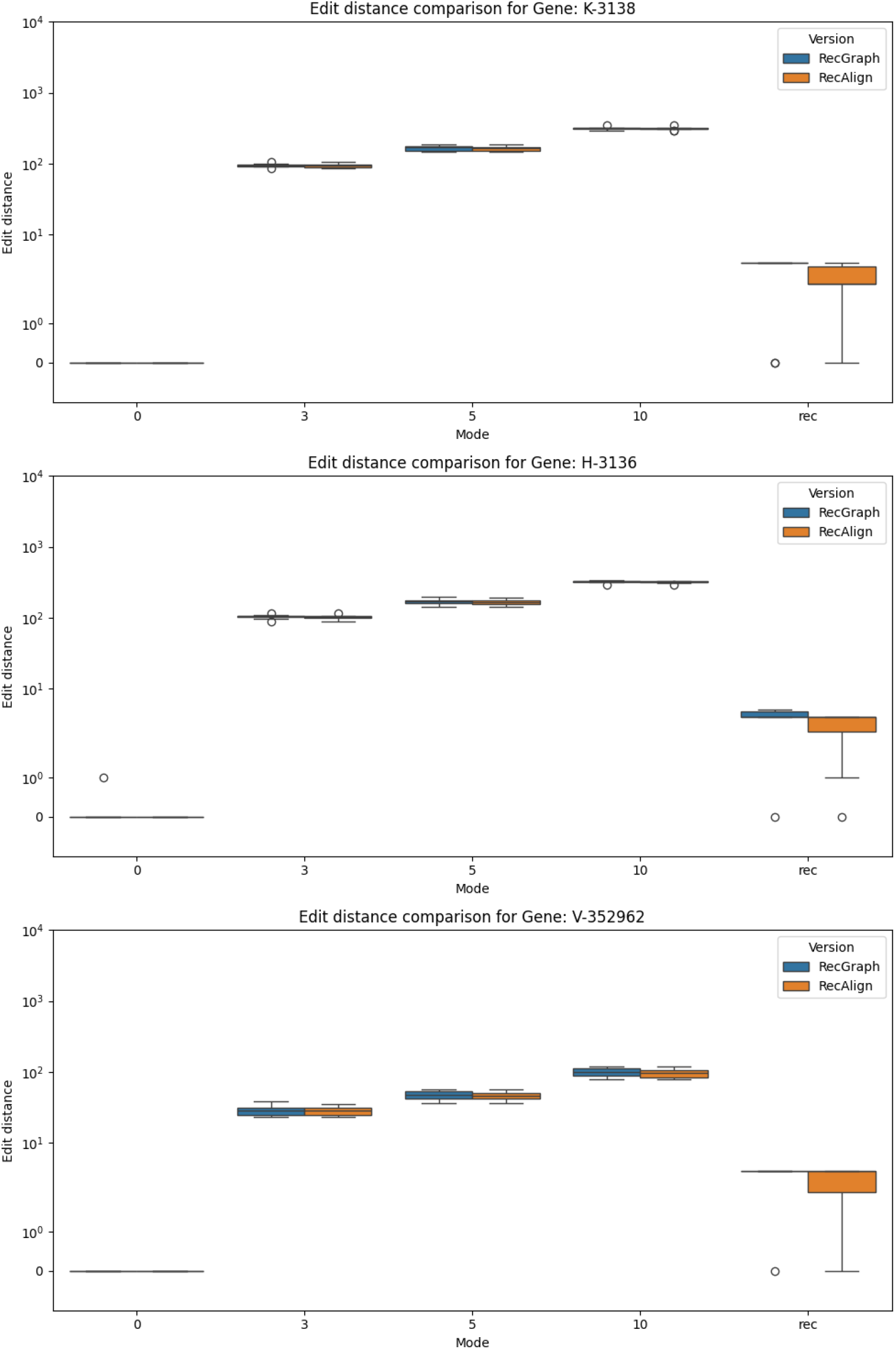
Edit distance distribution of RecGraph(blue) and RecAlign (orange) at different error rates and recombinations

**Figure 2:**
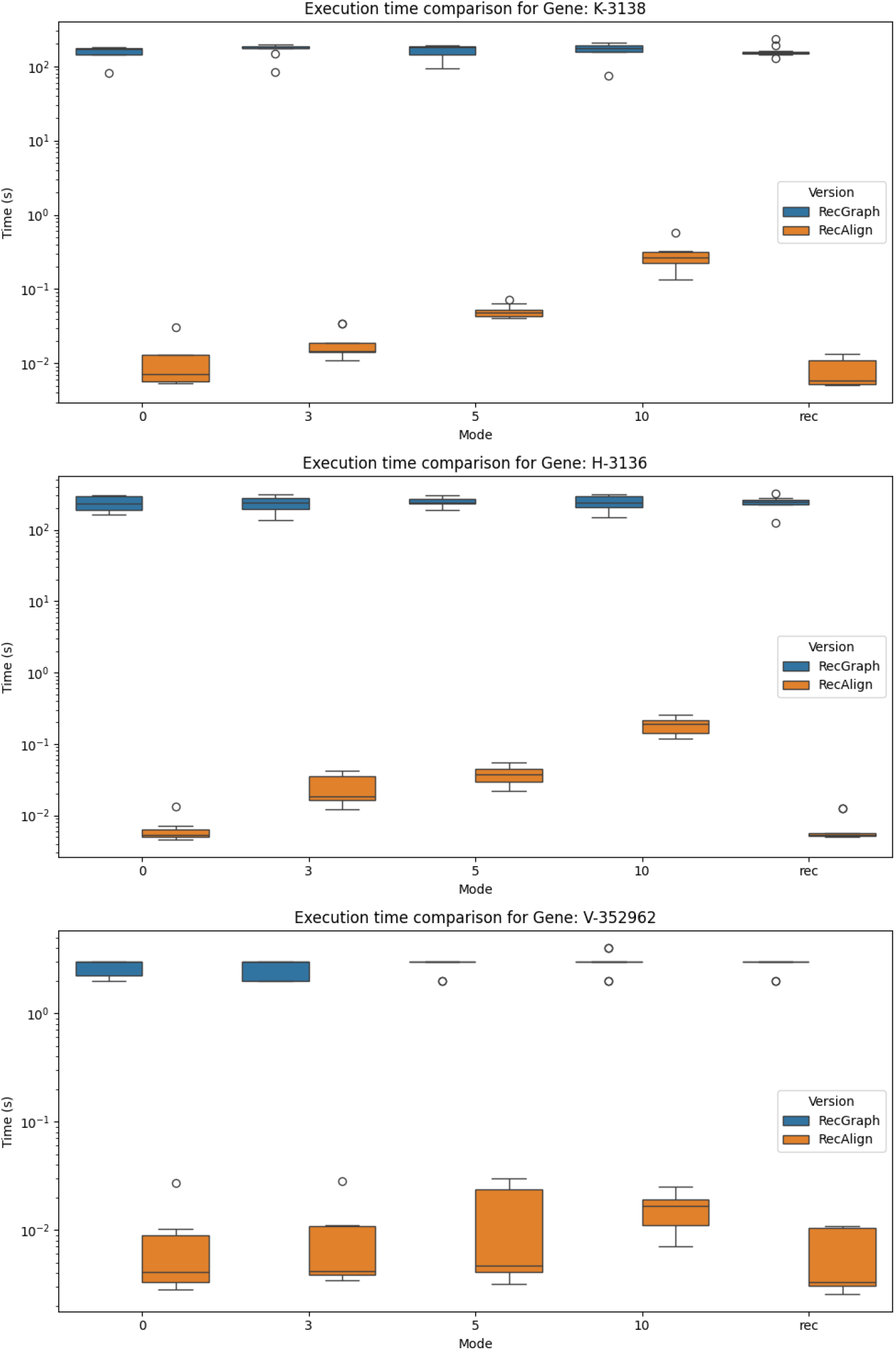
Execution time distribution of RecGraph(blue) and RecAlign (orange) at different error rates and recombinations

**Figure 3:**
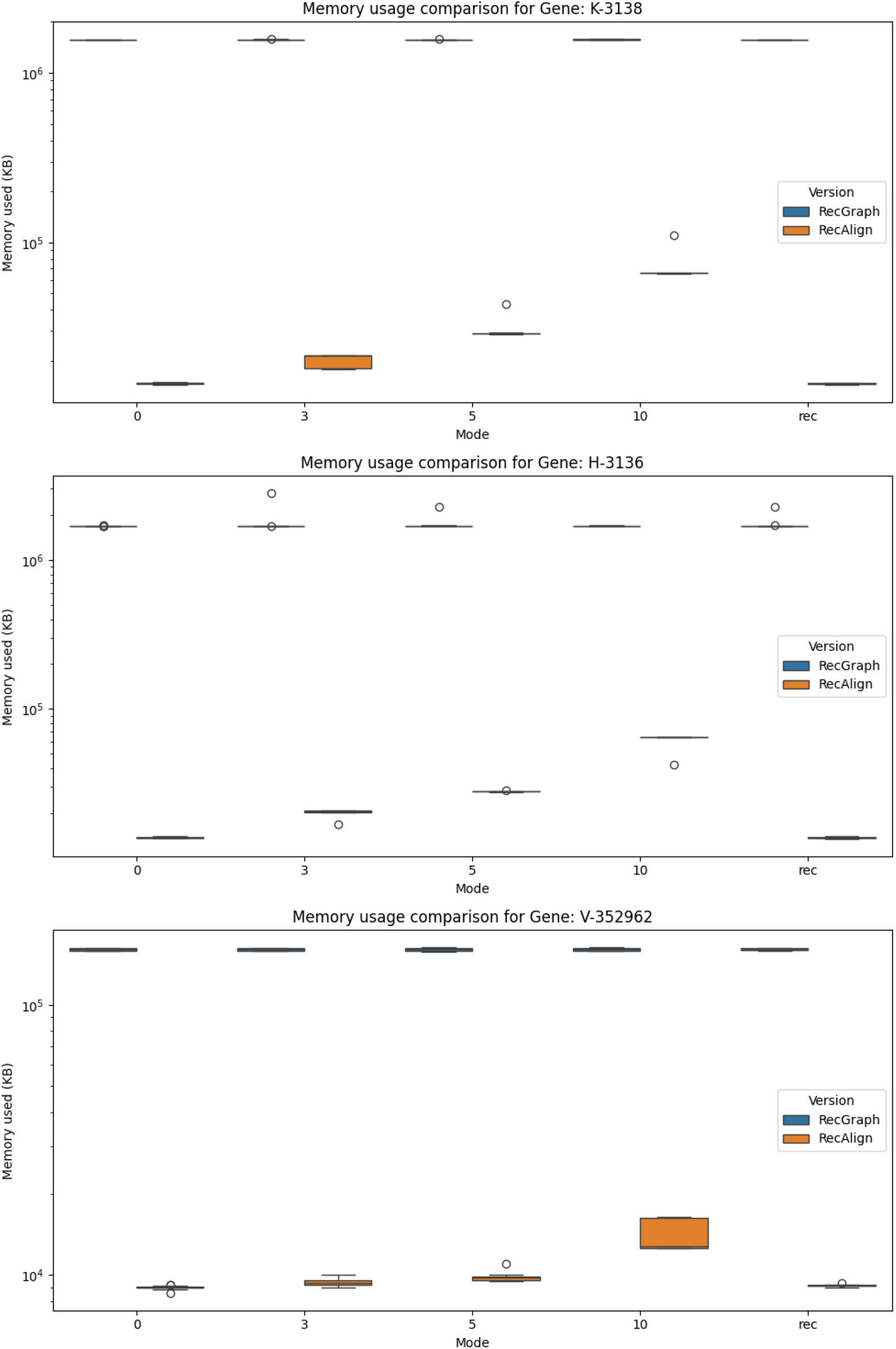
Peak memory distribution of RecGraph(blue) and RecAlign (orange) at different error rates and recombinations

#### 4.2.2 RecAlign’s performances analysis

In this second part of the experiment, we tried to focus more on what is affecting the performances of RecAlign, either the error rate or the number of recombinations, in order to confirm the preliminaries results from the previous experiment. We decided to simulate all combination of errors (0,3,5%) and recombinations (0,1,2), and then align every sequence against its respective graph.

We aggregate all results, computing a mean score for both memory and time. We then decided to keep track of the percentage of vertices of the alignment graph that are actually visited by the procedure. Note that this value can also be seen as the percentage of cells of the dynamic programming matrix computed.

We firstly set the maximum number of recombinations allowed in the alignment to the exact number of recombinations present in the read (with the parameter kset to 0 if no recombination was present in the read, 1 if one was present or 2 when there were two recombinations). The results are reported in figure 4. Firstly, we can see how all three parameters measured are strongly linked together, with a similar trend in all plots. From the experiment, however, emerged that, while for an error rate close to 0 the number of recombinations is not influencing the performances, as the error increases, also the effect of the maximum number of recombinations allowed increases, contradicting what we suspected after the previous experiment.

**Figure 4:**
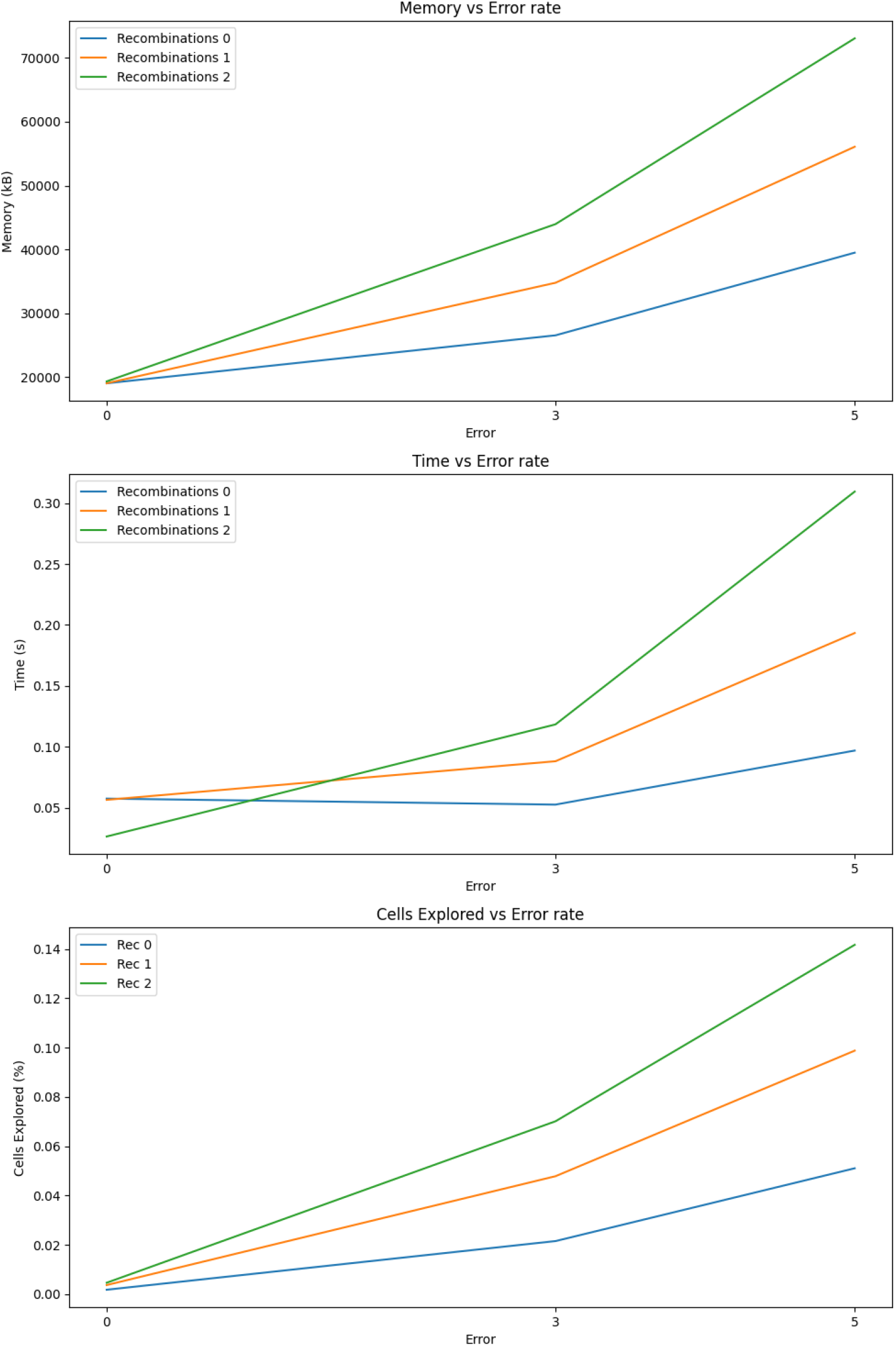
Mean execution times, peak memory used and percentage of nodes explored of RecAlign on the complete set of HLA genes, with different error rates and recombinations (variable k).

We then decided to repeat the experiment, setting the maximum number of recombinations allowed to 3 for every read, without considering the actual number of recombination present in it. Please note that the maximum number of recombination allowed implies that it is possible to have *up to k* recombinations, it does not require having exactly *k* recombinations. The results are shown in figure 5. It emerges that, in this situation, the number of recombinations seems to have no impact on the computation costs, while the error rate seems to have a similar impact. We can then assume that it is not the actual number of recombinations present in the read affecting RecAlign’s performances, but the maximum number of recombinations allowed in the alignment. That is due to the fact that adding more opportunity of introducing recombination events leads to a higher exploration of the possible solution’s space.

**Figure 5:**
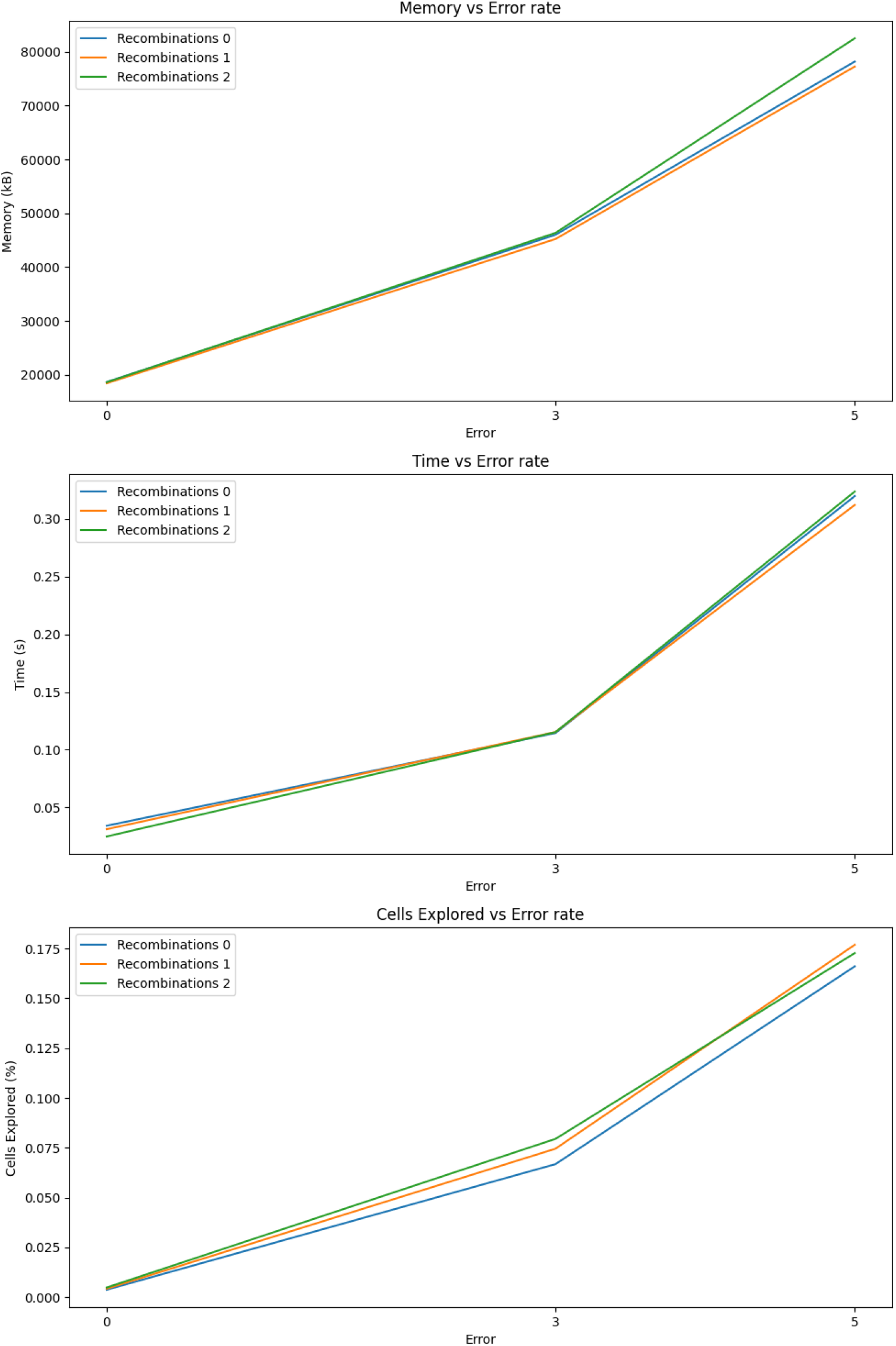
Mean execution times, peak memory used and percentage of nodes explored of RecAlign on the complete set of HLA genes, with different error rates and recombinations (fixed k).

One last consideration regards the *k* size of the seeds. During all the experiments, we chose to set it to 8, in order to balance the time needed to build the *h* score and to explore the alignment graph. A larger value for *k* tends to increase the time of exploration, while reducing the time needed for heuristic computation, the opposite for smaller values. However, this value is also linked to the error rate presented by the sequence. Sequences with higher error rates tend to need a lower *k* size in order to compute a better *h* score, because the probability of finding a large match decreases. The opposite happens when the error rate is low and a large *k* allows exploiting more the *skip ahead* step described in 3.3.

#### 4.2.3 Comparison with other tools

In this final experiment, we compare RecAlign with some other state-of-the-art aligners on the full set of HLA genes. For each gene we extracted the haplotypes present in the pangenome graph. In order to simulate recombination events, we then created new haplotypes by combining two or three original haplotypes, by choosing a random recombination point. We have generated 10 recombinant haplotypes with 1 recombination and 10 recombinant haplotypes with 2 recombinations for each gene. For some genes, we could not actually create all those recombinant haplotypes, since the number of original haplotypes was too small — in fact, we do not allow two recombinants from the same pair of original haplotypes. From each recombinant haplotype, we have simulated reads by adding random noise, that is changing, inserting, or deleting single nucleotides, with a 0, 3% or 5% error rates. For each gene and each combination of parameter values (error rate and number of recombinations), we have simulated one read.

To estimate the alignment quality we employ the alignment cost, where each edit operation costs 1 and each recombination costs 4. Accordingly, RecAlign computes a global alignment, where we allow at most 2 recombinations, with a recombination penalty set to 4. The results are shown in Table 1.

**Table 1:** Mean edit scores and not aligned reads for each gene across tools.

| Gene | GraphAligner |  | RecAlign |  | Minichain |  |
| --- | --- | --- | --- | --- | --- | --- |
|  | Mean Cost | Not Aligned | Mean Cost | Not Aligned | Mean Cost | Not Aligned |
| B-3106 | 109.8 | 0/87 | 91.26 | 0/87 | 152.31 | 0/87 |
| C-3107 | 2631.74 | **4/90** | 92.27 | 0/90 | 147.11 | 0/90 |
| DMA-3108 | 122.17 | 0/83 | 116.63 | 0/83 | 126.88 | 0/83 |
| DPB1-3115 | 403.6 | 0/93 | 368.8 | 0/93 | 428.61 | 0/93 |
| DRB4-3126 | 342.04 | 0/75 | 335.21 | 0/75 | 359.67 | 0/75 |
| E-3133 | 129.78 | 0/87 | 128.78 | 0/87 | 133.31 | 0/87 |
| G-3135 | 116.94 | 0/93 | 110.39 | 0/93 | 123.78 | 0/93 |
| J-3137 | 111.16 | 0/90 | 107.18 | 0/90 | 121.63 | 0/90 |
| K-3138 | 99.08 | 0/77 | 85.68 | 0/77 | 112.97 | 0/77 |
| L-3139 | 205.35 | 0/84 | 196.01 | 0/84 | 225.93 | 0/84 |
| MICA-100507436 | 389.86 | 0/64 | 373.75 | 0/64 | 414.09 | 0/64 |
| TAP1-6890 | 2812.71 | **76/93** | 231.69 | 0/93 | 243.43 | 0/93 |
| TAP2-6891 | 454.38 | **1/93** | 448.12 | 0/93 | 476.22 | **3/93** |
| V-352962 | 31.69 | 0/90 | 28.07 | 0/90 | 59.47 | 0/90 |

We can observe that RecAlign was able to compute alignments with lower costs for every single gene. In particular, GraphAligner was not able to produce any alignment for 3 reads belonging to gene C-3107, 1 belongings to gene TAP2-6891 and most of the ones belonging to TAP1-6890. Minichain instead was not able to align 3 reads of gene TAP2-6891.

## 5 Conclusions and future directions

We have proposed an A* approach to a recombination haplotype-aware sequence to graph alignment. This is the first exact algorithm that bounds the number of allowed recombinations and works on general variation graphs with cycles and inversions. The tool RecAlign that implements this approach largely improves RecGraphby decreasing the running time and the memory usage by at least 2 orders of magnitude. This allows us to map efficiently long reads to HLA gene pangenome graphs. Future work will be devoted to combine RecAlign with another aligner based on the seed-and-extend paradigm, such as giraffe [S^+^21] used in [GSN^+^18], allowing the latter to also incorporate recombinations. While the present work analyzes the use of a recombination haplotype-aware approach over human HLA genes and bacterial genomes, showing that it improves over the edit distance, we believe that a tool like RecAlign can highlight a biological interpretation of an alignment with multiple recombinations that was not previously possible due to the absence of such a procedure. Mainly, we plan to intensively apply RecAlign to more general graph representations that allow cycles and inversions as those constructed with [GG23].

## Footnotes

1 https://github.com/baku4/lt-fm-index

## References

[ACBC^+^24] Jorge Avila Cartes, Paola Bonizzoni, Simone Ciccolella, Gianluca Della Vedova, Luca Denti, Xavier Didelot, Davide Cesare Monti, and Yuri Pirola. RecGraph: recombination-aware alignment of sequences to variation graphs. Bioinformatics, 40(5):btae292, 04 2024.

[AW21] Tim Anderson and Travis J. Wheeler. An optimized FM-index library for nucleotide and amino acid search. Algorithms for Molecular Biology, 16(1):25, December 2021.

[B^+^22] Jasmijn A. Baaijens et al. Computational graph pangenomics: a tutorial on data structures and their applications. Natural Computing, 21(1):81–108, 2022.

[CGJ24] Ghanshyam Chandra, Daniel Gibney, and Chirag Jain. Haplotype-aware sequence alignment to pangenome graphs. Genome Research, page gr.279143.124, July 2024. Company: Cold Spring Harbor Laboratory Press Distributor: Cold Spring Harbor Laboratory Press Institution: Cold Spring Harbor Laboratory Press Label: Cold Spring Harbor Laboratory Press Publisher: Cold Spring Harbor Lab.

[DDA^+^13] Kate E. Dingle, Xavier Didelot, M. Azim Ansari, David W. Eyre, Alison Vaughan, David Griffiths, Camilla L. C. Ip, Elizabeth M. Batty, Tanya Golubchik, Rory Bowden, Keith A. Jolley, Derek W. Hood, Warren N. Fawley, A. Sarah Walker, Timothy E. Peto, Mark H. Wilcox, and Derrick W. Crook. Recombinational Switching of the Clostridium difficile S-Layer and a Novel Glycosylation Gene Cluster Revealed by Large-Scale Whole-Genome Sequencing. The Journal of Infectious Diseases, 207(4):675–686, February 2013.

[Dij59] E. W. Dijkstra. A note on two problems in connexion with graphs. Numerische Mathematik, 1(1):269–271, December 1959.

[DM10] Xavier Didelot and Martin CJ Maiden. Impact of recombination on bacterial evolution. Trends Microbiol, 18(7):315–322, 2010.

[EMTG23] Massimo Equi, Veli Mäkinen, Alexandru I. Tomescu, and Roberto Grossi. On the Complexity of String Matching for Graphs. ACM Transactions on Algorithms, 19(3):21:1–21:25, April 2023.

[Fre75] Michael L Fredman. On computing the length of longest increasing subsequences. Discrete Mathematics, 11(1):29– 35, 1975.

[GG23] Erik Garrison and Andrea Guarracino. Unbiased pangenome graphs. Bioinformatics, 39(1):btac743, 2023.

[GKI24] Ragnar Groot Koerkamp and Pesho Ivanov. Exact global alignment using A* with chaining seed heuristic and match pruning. Bioinformatics, 40(3):btae032, 01 2024.

[GSN^+^18] Erik Garrison, Jouni Sirèn, Adam M Novak, Glenn Hickey, Jordan M Eizenga, Eric T Dawson, William Jones, Shilpa Garg, Charles Markello, Michael F Lin, et al. Variation graph toolkit improves read mapping by representing genetic variation in the reference. Nature biotechnology, 36(9):875–879, 2018.

[Gus97] Dan Gusfield. Algorithms on Strings, Trees and Sequences: Computer Science and Computational Biology. Cambridge University Press, Cambridge, 1997.

[HNR68] Peter E Hart, Nils J Nilsson, and Bertram Raphael. A formal basis for the heuristic determination of minimum cost paths. IEEE transactions on Systems Science and Cybernetics, 4(2):100–107, 1968.

[IBV22] Pesho Ivanov, Benjamin Bichsel, and Martin Vechev. Fast and optimal sequence-to-graph alignment guided by seeds. In International Conference on Research in Computational Molecular Biology, pages 306–325. Springer, 2022.

[LAE^+^23] Wen-Wei Liao, Mobin Asri, Jana Ebler, Daniel Doerr, Marina Haukness, Glenn Hickey, Shuangjia Lu, Julian K. Lucas, Jean Monlong, Haley J. Abel, Silvia Buonaiuto, Xian H. Chang, Haoyu Cheng, Justin Chu, Vincenza Colonna, Jordan M. Eizenga, Xiaowen Feng, Christian Fischer, Robert S. Fulton, Shilpa Garg, Cristian Groza, Andrea Guarracino, William T. Harvey, Simon Heumos, Kerstin Howe, Miten Jain, Tsung-Yu Lu, Charles Markello, Fergal J. Martin, Matthew W. Mitchell, Katherine M. Munson, Moses Njagi Mwaniki, Adam M. Novak, Hugh E. Olsen, Trevor Pesout, David Porubsky, Pjotr Prins, Jonas A. Sibbesen, Jouni Sirèn, Chad Tomlinson, Flavia Villani, Mitchell R. Vollger, Lucinda L. Antonacci-Fulton, Gunjan Baid, Carl A. Baker, Anastasiya Belyaeva, Konstantinos Billis, Andrew Carroll, Pi-Chuan Chang, Sarah Cody, Daniel E. Cook, Robert M. Cook-Deegan, Omar E. Cornejo, Mark Diekhans, Peter Ebert, Susan Fairley, Olivier Fedrigo, Adam L. Felsenfeld, Giulio Formenti, Adam Frankish, Yan Gao, Nanibaa’ A. Garrison, Carlos Garcia Giron, Richard E. Green, Leanne Haggerty, Kendra Hoekzema, Thibaut Hourlier, Hanlee P. Ji, Eimear E. Kenny, Barbara A. Koenig, Alexey Kolesnikov, Jan O. Korbel, Jennifer Kordosky, Sergey Koren, HoJoon Lee, Alexandra P. Lewis, Hugo Magalhães, Santiago Marco-Sola, Pierre Marijon, Ann McCartney, Jennifer McDaniel, Jacquelyn Mountcastle, Maria Nattestad, Sergey Nurk, Nathan D. Olson, Alice B. Popejoy, Daniela Puiu, Mikko Rautiainen, Allison A. Regier, Arang Rhie, Samuel Sacco, Ashley D. Sanders, Valerie A. Schneider, Baergen I. Schultz, Kishwar Shafin, Michael W. Smith, Heidi J. Sofia, Ahmad N. Abou Tayoun, Françcoise Thibaud-Nissen, Francesca Floriana Tricomi, Justin Wagner, Brian Walenz, Jonathan M. D. Wood, Aleksey V. Zimin, Guillaume Bourque, Mark J. P. Chaisson, Paul Flicek, Adam M. Phillippy, Justin M. Zook, Evan E. Eichler, David Haussler, Ting Wang, Erich D. Jarvis, Karen H. Miga, Erik Garrison, Tobias Marschall, Ira M. Hall, Heng Li, and Benedict Paten. A draft human pangenome reference. Nature, 617(7960):312–324, May 2023. Number: 7960 Publisher: Nature Publishing Group.

[LGS02] Christopher Lee, Catherine Grasso, and Mark F. Sharlow. Multiple sequence alignment using partial order graphs. Bioinformatics, 18(3):452–464, 2002.

[RM20] Mikko Rautiainen and Tobias Marschall. Graphaligner: rapid and versatile sequence-to-graph alignment. Genome biology, 21(1):253, 2020.

[S^+^21] Jouni Sirèn et al. Genotyping common, large structural variations in 5,202 genomes using pangenomes, the Giraffe mapper, and the vg toolkit. bioRxiv:2020.12.04.412486, 2021.

[SGN^+^19] Jouni Sirèn, Erik Garrison, Adam M. Novak, Benedict Paten, and Richard Durbin. Haplotype-aware graph indexes. Bioinformatics, 2019.

[SRS02] Rainer Spang, Marc Rehmsmeier, and Jens Stoye. A novel approach to remote homology detection: jumping alignments. J. Comput. Biol., 9(5):747–760, 2002.

